# Marine Cyanobacteria Tune Energy Transfer Efficiency in their Light-harvesting Antennae by Modifying Pigment Coupling

**DOI:** 10.1101/656173

**Authors:** Yuval Kolodny, Hagit Zer, Mor Propper, Shira Yochelis, Yossi Paltiel, Nir Keren

## Abstract

Photosynthetic organisms regulate energy transfer to fit to changes in environmental conditions. The biophysical principles underlying the flexibility and efficiency of energy transfer in the light-harvesting process are still not fully understood. Here we examine how energy transfer is regulated *in-vivo*. We compare different acclimation states of the photosynthetic apparatus in a marine cyanobacterial species that is well adapted to vertical mixing of the ocean water column and identify a novel acclimation strategy for photosynthetic life under low light intensities. Antennae rods extend, as expected, increasing light absorption. Surprisingly, in contrast to what was known for plants and predicted by classic calculations, these longer rods transfer energy faster *i.e.* more efficiently. The fluorescence lifetime and emission spectra dependence on temperature, at the range of 4-300K, suggests that energy transfer efficiency is tuned by modifying the energetic coupling strength between antennae pigments.

## introduction

Photosynthesis is regarded as one of the most efficient energy transduction processes in nature. Although it rarely operates at maximum efficiency, the high quantum efficiency generates a large dynamic range that allows the regulation of energy fluxes through the photosynthetic apparatus. This remarkable flexibility of the photosynthetic process is critical under changing environmental conditions ^1,2^.

The physical mechanisms by which the energy migration is controlled in the photosynthetic apparatus is under intense investigation. Traditionally, these mechanisms have been semi-classically modeled as dipole-dipole interactions between adjacent chromophores using Forster resonance energy transfer (FRET) amidst an incoherent bath ^3^. Evidence of long-lived quantum coherence, measured by 2D photon-echo spectroscopy, raised the possibility that quantum phenomena may play a role in the energy transfer process ^4,5^. The nature of these coherences, whether electronic or vibronic, is still under debate ^6–9^. Theoretical calculations demonstrate that both purely classic and purely quantum exciton energy transfer (EET) through antenna systems have their limitations. However, a combination of quantum and classic processes could enhance EET efficiency ^3^. The suggested combined mechanisms envision short-lived coherent domains encompassing small subsets of antenna pigments. It implies that at the border between classic and quantum regimes, small structural changes could have large effects on energy transfer ^10^.

It is well established that the sizes of photosynthetic units can reach hundreds of pigments. A recent structural study resolved an intact phycobilisome (PBS) antenna with thousands of pigments ^11^. Elucidating energy transfer dynamics in such large-scale systems will benefit from studies of intact systems, within their biological context. However, the study of ultrafast processes by advanced spectroscopy is usually done on isolated pigment-protein complexes. The benefits of working with small numbers of pigments are clear, but the biological relevance of such experiments is limited by their small scale. *In vivo* measurements present more complex scenarios that cannot be fully captured in simplified isolates. As phrased by Aristotle, “which have several parts and in which the totality is not, as it were, a mere heap, but the whole is something beside the parts”^1^. An example for this approach is the study of desiccation tolerant desert crust cyanobacteria ^12^. That study demonstrated the link between small changes in PBS structure and large quenching effects in the PBS antenna directly, under a desiccation/rehydration scenario where water availability triggers quick energy transfer tuning. Since in desert environments light intensities are high, it is interesting to also examine organisms that are adapted to life where light is a limiting factor.

In this study we looked for an organism which is adapted to an environment where (a) light can be limiting to the extreme, and (b) The intensity and quality of light varies, requiring acclimation of photosynthetic systems to a broad range of light regimes. Marine *Synechococcus* spp. are obvious candidates for such a study. The marine environment provides a matrix of light and environmental conditions that are even more complex than those in a terrestrial desert. In the marine environment, where as much as 50% of the world’s primary productivity takes place ^13^, light is the major factor limiting the abundance of photosynthetic organisms. Light intensity attenuates exponentially as it penetrates deeper into the water column and its spectrum is narrowed to blue (∼490 nm) ^13,14^.

An important perturbation of the open oceans water column is vertical mixing, which greatly affects the dynamics of *Synechococcus* populations ^15^. Mixing of a stratified water column, typically to a depth of a few hundred meters, homogenously distributes the nutrients and the planktonic organisms along the water column. The only resource that cannot be “mixed” is sunlight. As a result, an organism that lives near the surface with plenty of available light, may rather abruptly find itself with very little light if it reaches the deeper layers ^16^. The response to varying light conditions (photon flux density and spectral distribution), triggers a cascade of photoacclimation responses affecting the form and function of light harvesting systems ^17–19^.

*Synechococcus* are small unicellular coccoid cyanobacteria responsible for an estimated 25% of global primary productivity. They use phycobilisomes (PBSs) as their light-harvesting antenna. Among them, *Synechococcus* WH8102 represents an abundant clade that is well adapted to mixing regime and can be found throughout the water column ^15^. Studies of marine *Synechococcus*, including WH8102, demonstrated their ability to acclimate to the wide range of light intensities experienced along the water column. ^20^.

*Synechococcus* WH8102 PBS consist of phycobiliprotein (PBP) αβ dimers organized into trimer rings which, in turn, couple to create the hexamers which form the basic subunit of the core and stacked rods ^21^. The core is made of allophycocyanin (APC) and the rods of phycocyanin (PC) and phycoerythrin (PE). These PBPs consist of an apo-protein and contain 1-3 open-chain tetrapyrrole bilin chromophores. APC and PC both bind phycocyanobilin (PCB). The different protein environments generate a bathochromic shift in APC absorption ^22^, 620 nm in PC and 650 nm in APC. The rod extremities are constituted by two structurally different forms of PE: PEI and PEII. PEI binds either only phycoerythrobilin (PEB, absorption peak 545 nm), or PEB and phycourobilin (PUB, absorption peak 495 nm). PEII binds both PEB and PUB, but with fewer PEB than does PEI ^20^. *Synechococcus* demonstrates chromatic adaptation, able to modify the PEB into PUB and reversibly according to the available light spectrum ^23–25^.

Importantly, *Synechococcus* WH8102 has the highest reported content of phycourobilin, the pigment which best absorbs the blue light that penetrates ocean waters ^26,27^. The fact that it possesses particularly large PBSs challenges energy transfer processes, since energy transfer occurs over longer distances. According to classical mechanics approach, energy transfer efficiency is expected to decrease with a larger number of pigments and rod length. The optimal length was calculated to be very short, on the order of a few trimers ^28^.

Photo-physiological studies conducted on the response of marine *Synechococcus* species to light intensity gradients ^20^ demonstrated a large dynamic range of photosynthetic activity. In the study we present here, we take advantage of these photo-acclimation processes, to study how photosynthetic units tune their energy transfer rates. Using *Synechococcus* WH8102, we generated a testing ground for studying ultrafast energy transfer processes in large antennae structures with complex pigment compositions *in vivo*. Our analysis includes physiological, biochemical and spectroscopic measurements of energy transfer dynamics, conducted while comparing acclimated states of this marine cyanobacterium to different light regimes.

## Results and discussion

In order to compare the function of photosynthetic units within cells acclimated to light conditions representing shallower and deeper water column depths, we grew *Synechococcus* WH8102 cultures under blue LED light illumination, which resembles the light spectrum that penetrates the ocean (Supplementary fig.1). We compared cultures grown at different blue light intensities: 10 and 150 *µmol photons m*^−2^*s*^−1^, denoted “low light” and “medium light” respectively. It is important to note that in both cases the light intensity is lower than the intensity required for optimal growth, which is 207 *µmol photons m*^−2^*s*^−1 20^. Since light-induced stress was avoided, nonphotochemical quenching or photo-inhibitory losses of absorbed light energy are not expected under these growth conditions ^29^.

### Growth and primary productivity parameters

#### Synechococcus

WH8102 underwent photo-acclimation when grown under the two different blue light intensities. They exhibited extensive morphological and physiological modifications, linked to changes in their photosynthetic units. Immediately noticeable to the naked eye, is a change in the cultures color (fig. 1A). The acclimation process is evident within two days, as seen by several physiological parameters (fig. 1B, Supplementary figures 2–4) and is completed in about 7-10 days. Maximal apparent photosynthetic yield, measured as Fv/Fm, shows that the low light culture exhibits significantly higher values compared to the medium light culture (Table. 1), indicating higher photosynthetic capacity at low light conditions in cells grown under low light.

**Figure 1.**
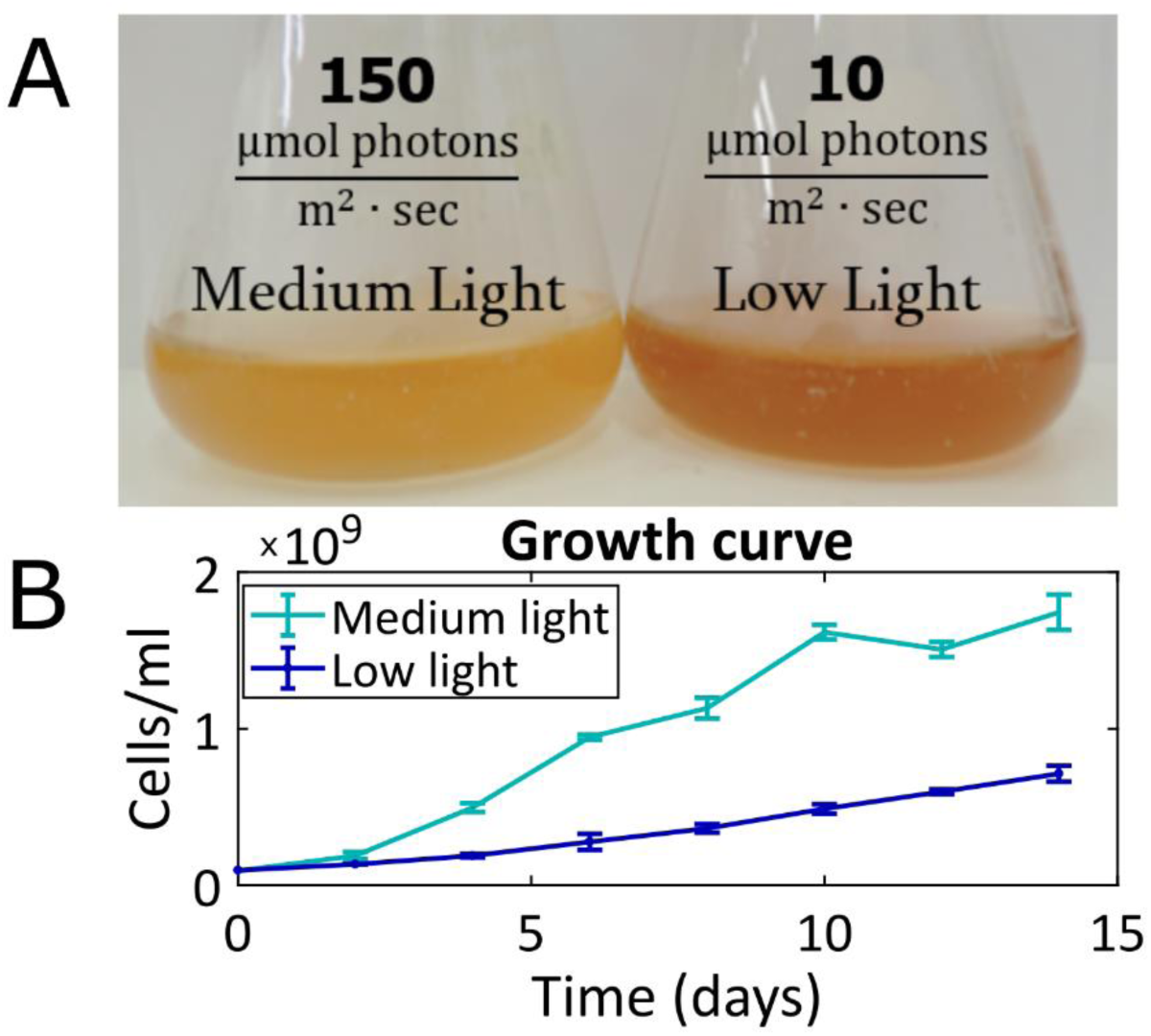
Photo-acclimation. (A) Synechococcus WH8102 cultures grown under two different blue light intensities for 5 days differ in color. (B) Growth curve under different light intensities, determined by FACS. While the color of low light cultures was stronger, medium light cultures exhibited higher cell densities. SD n=3; The experiments were independently repeated two additional times with comparable results.

**Table 1.** Differences in the photosynthetic units’ composition and function (n – number of independent repeats).

|  | Low Light | Medium Light | LL/ML ratio |
| --- | --- | --- | --- |
| **Chl/protein**<br>($\mu\text{g chl mg}^{-1}\text{protein}$) | 274 | 37 | 7.4 |
| **PUB/PC** | $11.9 \pm 1.5$ ($n = 4$) | $4.9 \pm 1.5$ ($n = 4$) | 2.4 |
| **PEB/PC** | $6.2 \pm 0.9$ ($n = 4$) | $2.9 \pm 0.8$ ($n = 4$) | 2.1 |
| **Fv/Fm** | $0.43 \pm 0.04$ ($n = 2$) | $0.22 \pm 0.03$ ($n = 2$) | 1.9 |

#### Synechococcus

WH8102 was able to grow under both low and medium light intensities. Cell numbers increased at a slower rate under low light. However, unexpectedly, the side scatter area parameter (SSC-A) of the fluorescence-activated cell sorting (FACS) measurement (Supplementary fig.3) raised the possibility that cell sizes were larger when grown under lower light. This prompted us to examine cellular morphology more closely.

### Cellular morphology

Confocal fluorescence microscopy and transmission electron microscopy (TEM) confirmed that cells change their shape, size and internal structure in response to the different blue light intensities. Under low light, the median size cell increased, and the size distribution was broader, as seen from the SSC-A statistics (fig. 2A) and the quantification of confocal images (Supplementary fig.2). In addition, TEM revealed that the cells changed their structure from round coccids under medium light to elongated under low light (fig. 2B). The elongated shape of the cells under low light, along with the broad size distribution, is consistent with their slower growth rate. Lastly, under medium light cells typically had one thylakoid membrane while under low light three to four membranes were observed, organized in uniformly distributed distances. Photosynthetic units are embedded in the thylakoid membranes, so an increase in the number of membranes indicates an increase in the number of photosynthetic complexes per cell. These results are similar to photo-acclimation changes observed in other marine cyanobacterial species ^30^.

**Figure 2.**
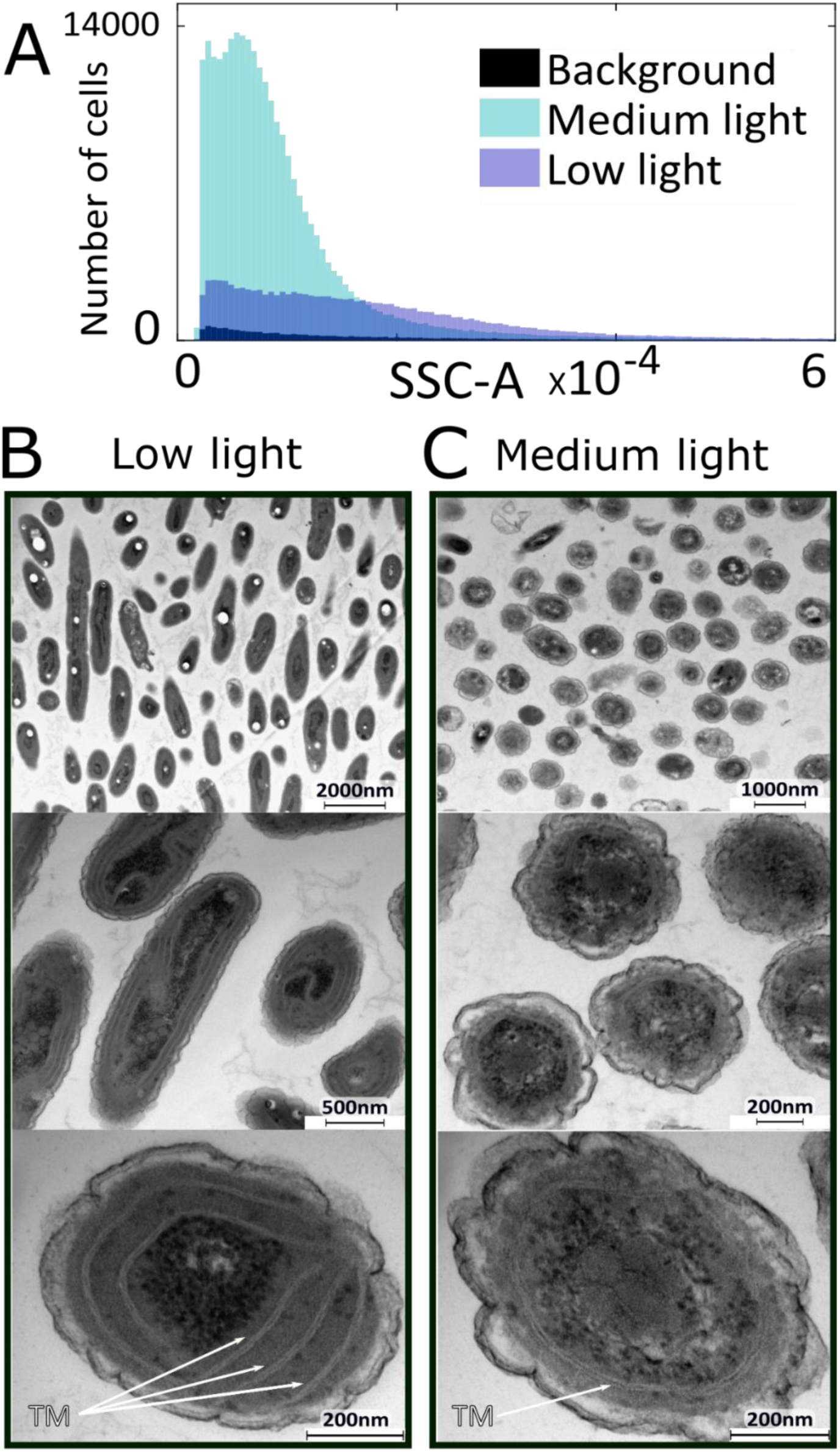
Cellular morphology. (A) Histogram of side scatter area (SSC-A) measured by FACS. Cultures grown under low light exhibited larger scattering values. Their mean size is about twice the size of cells grown under medium light and their scattering distribution is broader. These results indicate larger cell sizes. (B) Transmission electron microscopy of medium light cells. Typically, the cells are round, with one thylakoid membrane (TM) circling the cell. (C) Low light cells have longitude structure and contain about three thylakoid membranes. Additional information on cell sizes and architecture can be found in Supplementary fig.2.

The increased cellular surface area has the advantage of a higher light absorption cross-section, which is beneficial when light is limiting, provided that the energy transfer efficiency remains high. It is important to note that an increase in cross-section and cell size has the disadvantage of a reduced surface area per volume ratio, which may limit the assimilation of nutrients ^31^. The fact that nutrients are typically more abundant deeper within the ocean water column mitigates this risk. Under our experimental conditions, however, nutrients were abundant in both cultures.

### Photosynthetic complexes

As implied by the higher number of thylakoid membranes, under low light the number of photosynthetic units is higher. This is supported by the fact that chlorophyll *a* absorption, a component of both photosystems, was increased 8 folds at low light as compared to medium light on a per-cell basis (fig. 3). Similarly, on a protein basis, chlorophyll content increased 7.4 folds (Table 1). The ratio of PSI to PSII was slightly increased under low light, as demonstrated by protein immunoblotting (Supplementary fig.5). A prominent change is also evident in the composition of the photosynthetic antennae pigments. Examining the pigment content of the antenna, our results show that under low light, the PE content increases. The ratios between the different pigments change, as seen in table 1. Specifically, the ratios of PUB and PEB to PC absorption increase approximately two folds. These results are comparable to a previous study done on *Synechococcus* WH8102 ^20^.

**Figure 3.**
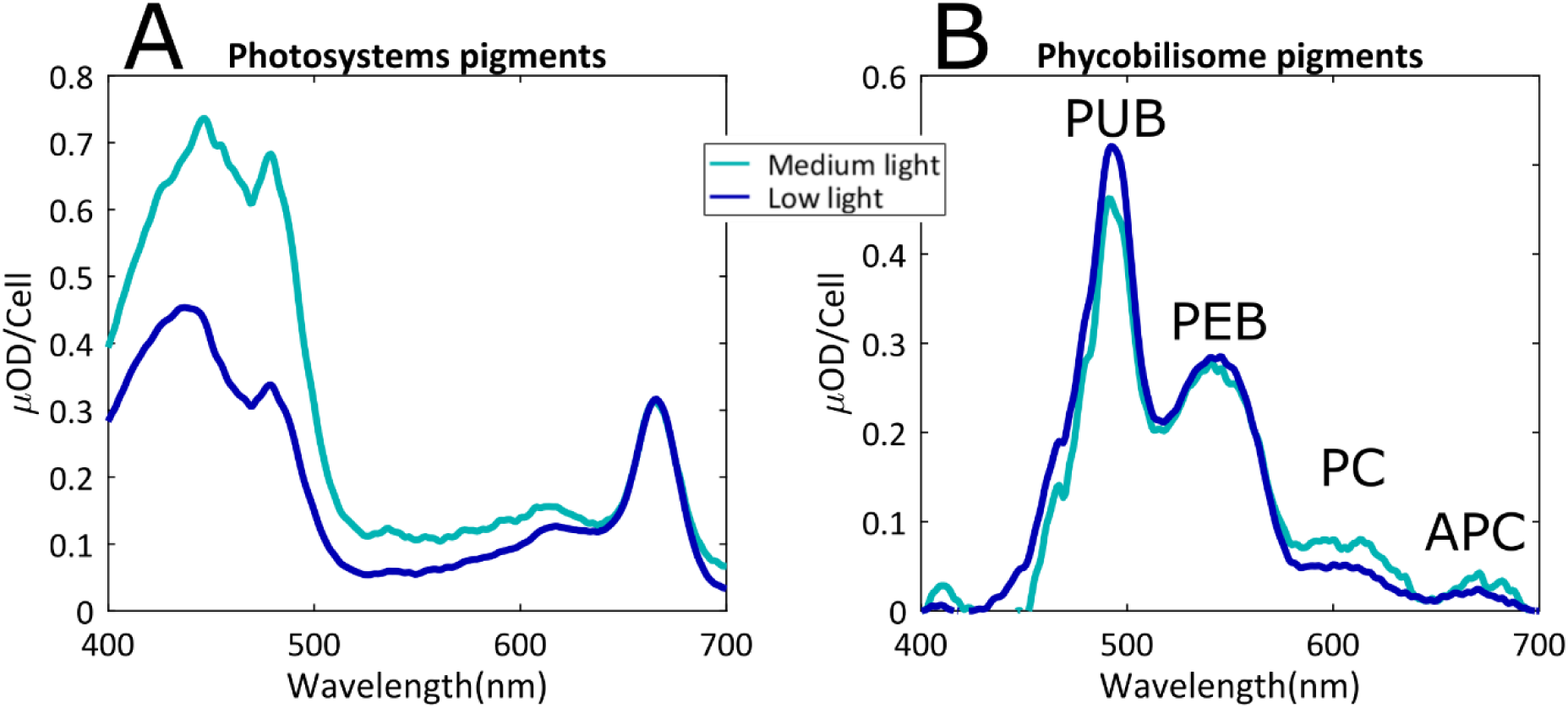
Photosynthetic complexes content. Absorption spectra of extracted pigments from the cells measured using integrating sphere. Values are normalized on a per cell basis. Low light cells contained considerably more pigments and their data was divided by eight to fit on the same scale. (A) Methanol extraction solubilizing hydrophobic chlorophylls and carotenoids. Since chlorophyll has two absorption peaks and the first excited state absorption is equal, the difference in in the absorption between 450-500nm is a result of a larger quantity of carotenoids in medium light. (B) MTN solubilized pigments showing the phycobilisome pigments (phycobilins). Lower light has higher PUB:PEB but lower PC:PEB ratio. In-vivo absorption spectra appears in Supplementary fig.6.

PBSs increase in size under low light ^32^. Absence of PC rod linker (LR) genes such as cpcC or cpcD in *Synechococcus* WH8102 suggests that under all light conditions there is only one hexameric PC disk per PBS rod ^32,33^. Taking this into account, it is most likely that under low light, where proportions of PUB and PEB to PC are higher, the rods become longer. According to Six and co-workers ^32^ PEII varies more than PEI, therefore it is reasonable to conclude that the number of PEII disks in each rod is higher under low light than under medium light. These considerations generated the model presented in fig. 4. Similar structures were suggested in studies of other *Synechococcus* specie^34^.

**Figure 4.**
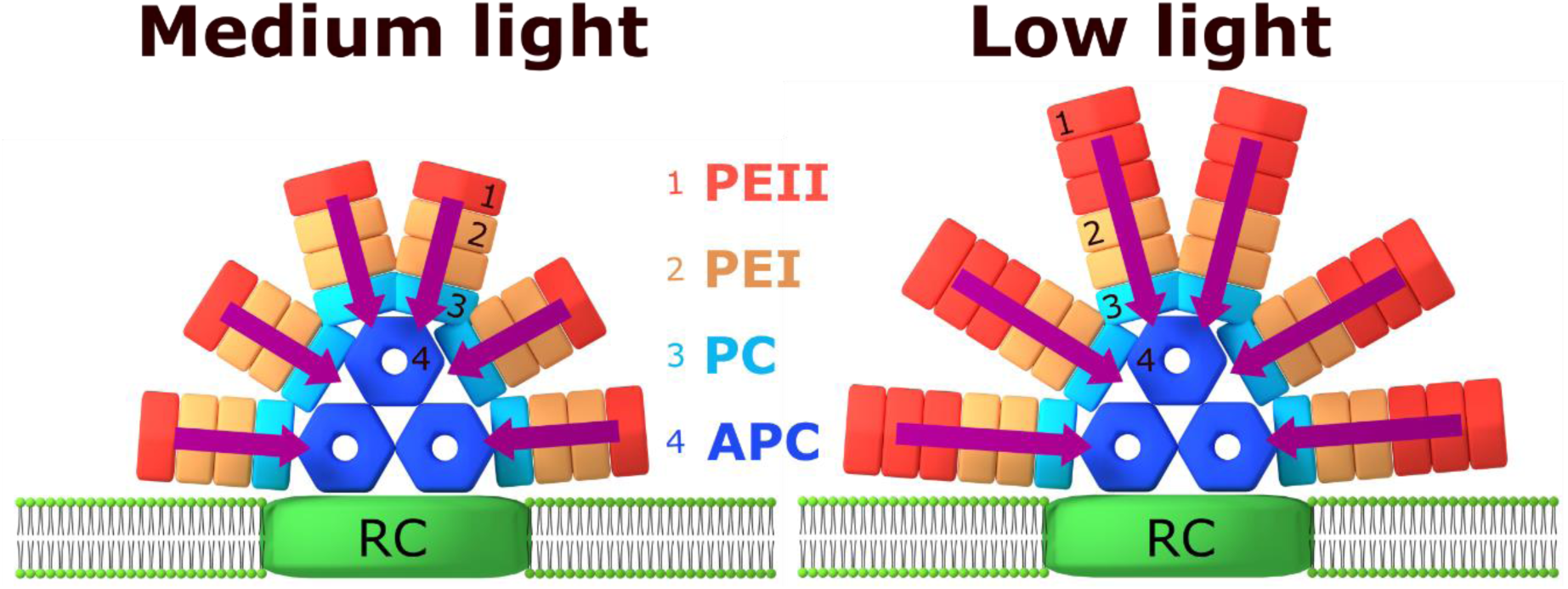
PBS structure under different light intensities. Suggested PBS scheme, showing the differences between the complexes grown under low versus medium light. Under low light the PBS is larger, with longer rods that have additional PEII hexamers (with higher PUB content). Energy absorbed at the extremities travels longer distance to the reaction centers (RCs). Structures are based on the pigment ratios found in this study as well as in Six and co-workers ^32^ and the structural organization of PBS in other strains that was published before ^34^, suggesting it is typically comprised of three APC cylinders and six rods.

These structures raise an interesting problem. While bigger light harvesting antennae containing additional PUB have an increased absorption cross-section, they generate a longer pathway for energy transfer to the reaction centers. Theoretical models using FRET demonstrated that an increase in rod length is expected to result in lower energy transfer efficiency ^28^. However, we found the functionality of each photosynthetic unit is improved under low light, as seen from the maximal apparent photosynthetic yield (Fv/Fm) values. To further probe the issue of energy transfer efficiency, we examined fluorescence spectra and lifetime characteristics.

### Energy transfer properties

Steady state fluorescence spectra of *Synechococcus* WH8102, at room temperature, allow us to compare the energy flow through the photosynthetic apparatus. Fig. 5A shows the emission spectra of cultures treated with DCMU, a PSII electron transfer inhibitor. It allows us to eliminate photochemistry and concentrate on exciton energy transfer to the reaction center. Excitation was set at 497 nm, the absorption peak of the PUB at the extremity of the rod. The spectra were normalized to the emission at 652 nm, the peak of the PC (assuming only one PC disk per rod). Interestingly, the emission in the spectral range of PC, APC and PSII is identical for both cultures (650-700 nm). This implies that the cells have maintained energy transfer homeostasis by regulating energy flow, so that the same flux that passes through the PC discs reaches the reaction center under both low and medium light.

**Figure 5.**
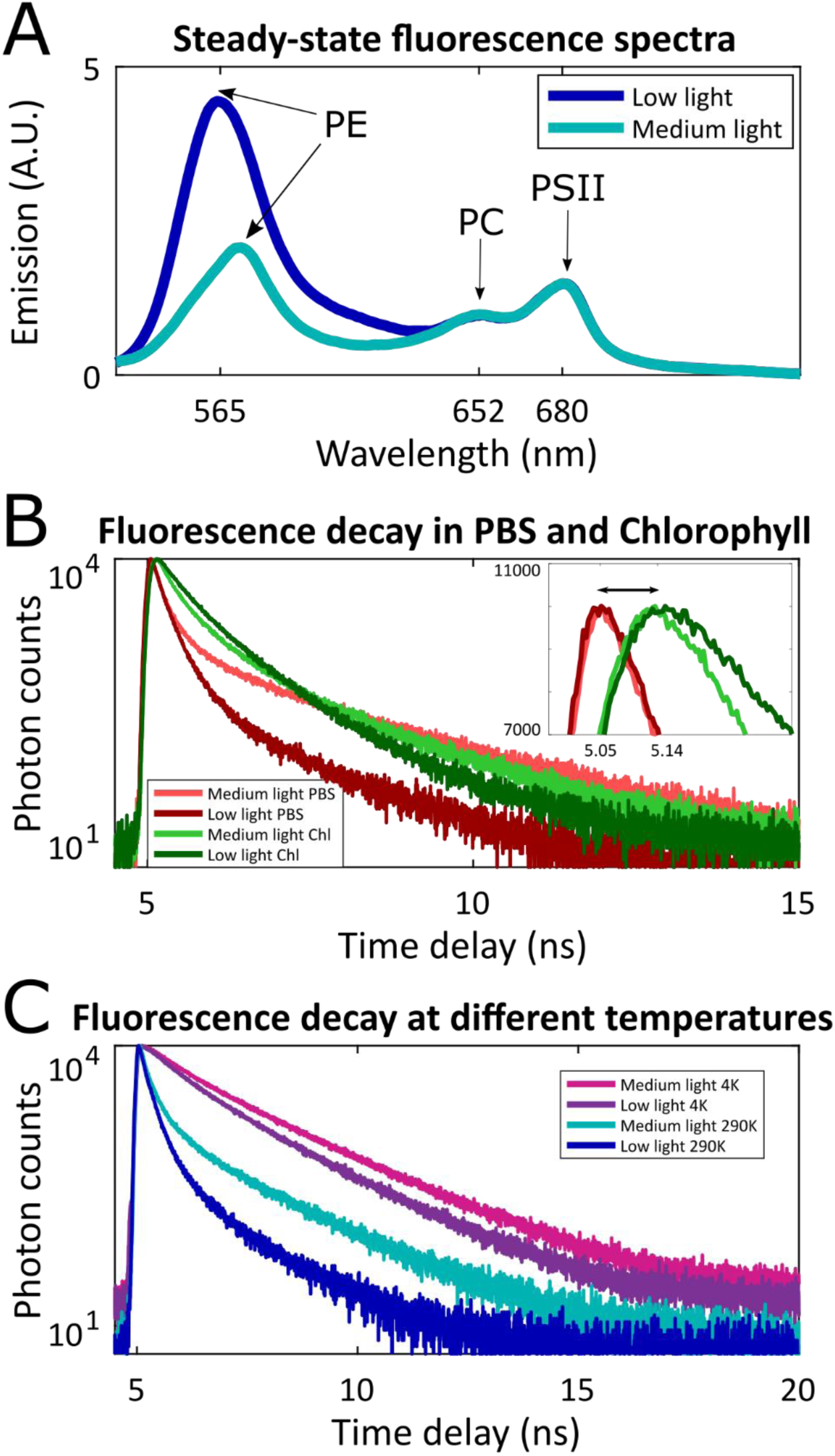
Energy transfer in photosynthetic apparatus. (A) in vivo emission spectra (excitation 497 nm). Emission was normalized to the peak of phycocyanin (652 nm). Samples treated with 5µM DCMU, to inhibit PSII photochemistry. PSI fluorescence is not observed at room temperature. The measurements were repeated in four independent experiments with comparable results. (B) Time resolved fluorescence spectroscopy in vivo (excitation at 485 nm, absorbed mainly by PE). Emission was collected in spectral windows of 500-675 nm (PBS window) or 675-750 nm (mainly chlorophyll, PS window). Insert shows the rise time of the chlorophyll peak. In both low and medium light, a 90 ps delay between the PS window and the PBS windows was observed. (C) Time resolved fluorescence at two temperatures (4K and 290K). Emission was integrated over both windows (500-750 nm).

The emission peak of the PE was shifted (565 nm at low light, 572 nm at medium light), consistent with the higher PUB:PEB ratio under low light (fig. 3). Moreover, PE emission was more than twice stronger in low light. This is expected considering the higher PE content. It may also imply that multiple PE units on the same rod cannot move energy simultaneously through the single PC disk, or that some of the PE pigments in cells grown under low light are not strongly coupled. A small number of uncoupled pigments can have a large contribution to the fluorescence yield.

To further investigate the rate of energy transfer following excitation of PUB, we performed measurements of the excited state’s lifetime. Time-correlated single photon counting approach (TCSPC) was used to measure the decay lifetime of the different components of the photosynthetic unit. Excitation was set at 485 nm targeting the extremities of the PBS rod, and emission was collected in three spectral windows, covering the phycobilins (PBS window, 500-675 nm), the APC terminal emitter and photosystem chlorophylls (PS window, 675 to 750 nm), and the overall system’s lifetime (Wide window, 500 to 750 nm). The measured lifetime, *τ*, is the typical lifetime of the system’s excited state, and is composed of different pathways which can be related to three major components: radiative (*τ*_*R*_), heat dissipation (*τ*_*k*_) and energy transfer (*τ*_*ET*_) pathways. Their combined effective value is formulated as a harmonic average 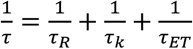

The PBS average lifetime was 0.12 ns under medium light, compared to 0.09 ns under low light, at room temperature (Fig. 5B). The fact that the lifetime is shortened under lower light is surprising. The opposite phenomenon was reported in plants, where lower light triggered an increase in lifetime ^35^. The shorter lifetime indicates that in each photosynthetic unit, the absorbed energy decays faster. This could either be due to higher dissipation of the energy, or due to faster energy transfer rate. The light intensity is low in both cases; therefore, multi-exciton processes are not relevant. Fv/Fm values are higher in low light acclimated cells, making disorder dissipation (energy quenching) by coupling to environment less probable ^10^. We hypothesized that higher energy transfer rates are responsible for the shorter life time.

The 90 ps shift between the peak of the PBS window trace and the PS window trace (fig. 5B insert) gives a rough estimation of the transfer rate between the phycobilisome and the reaction center. The fact that this timing is similar for both samples although the number of pigments per system is different supports the hypothesis that PBS energy transfer is accelerated under low light. The PS lifetime was 0.25 ns under medium light compared to 0.35 ns under low light. Examining the components of these decay curves indicates that the contribution of the fast decay component in the medium light is larger. This may be a result of the direct excitation of photosystem chlorophylls. While chlorophyll *a* absorption peak is around 430 nm, they do absorb, with lower efficiency, also at 497 nm (Fig. 3). Under medium light where chlorophyll absorption is not as masked by PBS absorption as in low light, the proportion of direct chlorophyll excitation will be higher.

As expected from the similar PC/PSII steady-state emission ratio but different PE/PC ratio seen in fig 5A, the major difference in the lifetime between the two samples is evident in the PBS fluorescence lifetime, suggesting this is the energy transfer step that is enhanced.

Fig. 5C shows that at 5K, where contributions of thermal vibrations are negligible, the lifetime at low light is still shorter. In fact, the difference in lifetimes increases with decreasing temperatures (Fig. 6A). Since radiation decay rates are intrinsic, it implies the energy transfer from the PBS rod extremity (the PUB) is indeed faster. This finding is surprising. FRET calculations predict a decrease with increasing number of pigments ^28^. Such a decrease was indeed demonstrated in plant systems, where longer lifetimes where reported in the large antenna of low light acclimated samples ^35^.

**Figure 6.**
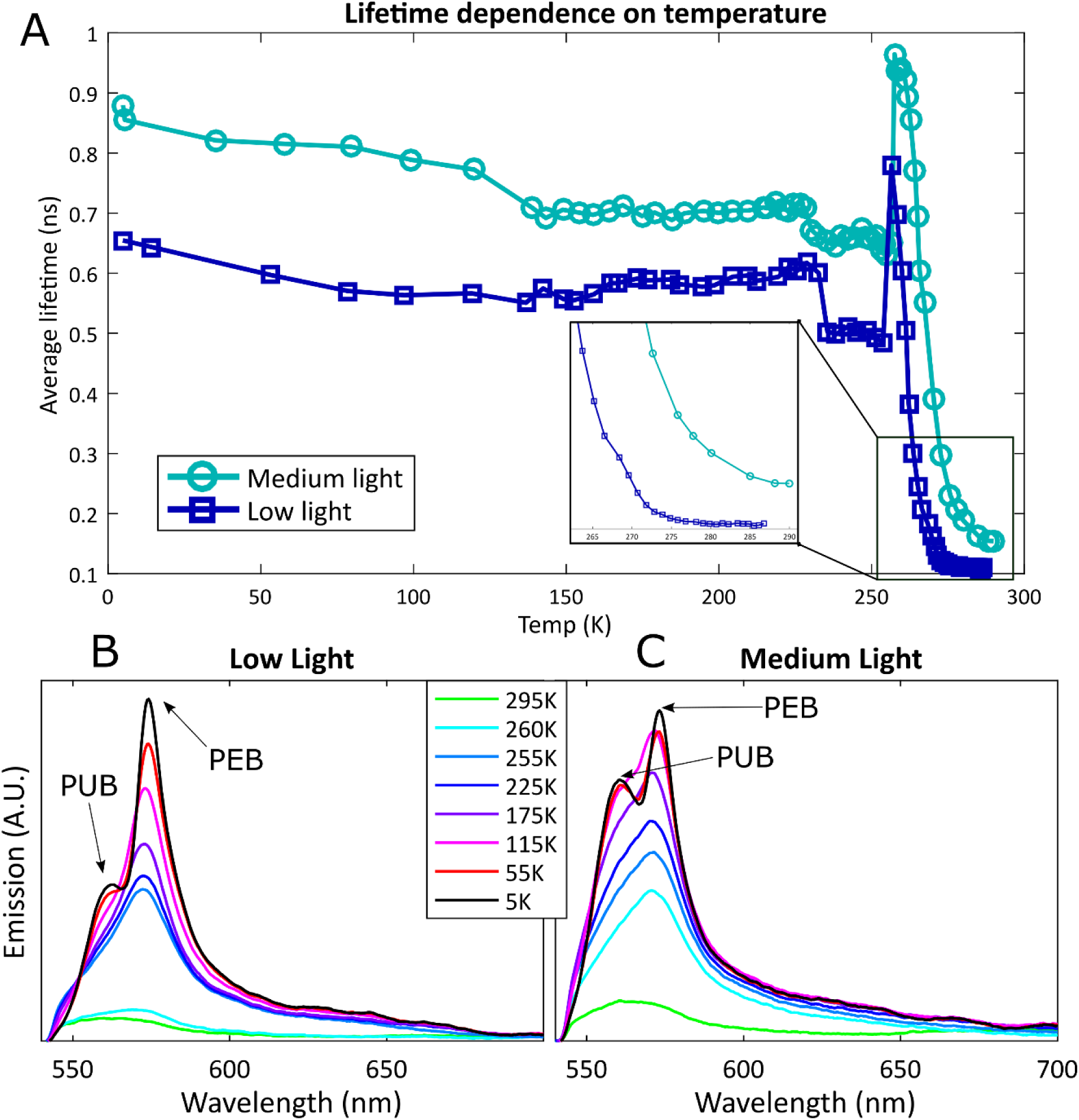
Temperature dependence. (A) Average lifetime of the photosynthetic complex as a function of temperature (measurement parameters as in 5C). (B,C) Fluorescence dependence on temperature. Spectrally resolved emission, following excitation at 450 nm, as a function of temperature from 295 down to 5K. Several representative temperatures are shown on graph.

### Pigment coupling

We hypothesized that faster energy transfer over a larger pigment system will require stronger energetic coupling between these pigments. To explore this hypothesis and inspect the coupling strength between the pigments, we measured the temperature dependence of both steady state and time-resolved fluorescence, from 300K down to 4K. Photochemistry is blocked below freezing at ∼273K, though the photosynthetic complexes remain intact. Spectral characteristics of a multi-chromophoric system are governed by the coupling between the chromophores, which, in turn, is affected by thermal vibrations. Hence, the temperature dependence of the lifetime and the emission spectrum can be used as a probe to study the coupling regime between pigments in the photosynthetic apparatus. This approach already produced interesting results *in vitro* ^36^, here we use it *in vivo.*

Fig. 6A shows the average fluorescence lifetime as a function of temperature. An interesting feature is the abrupt rise in lifetime at the temperature range of 250-265K. This phenomena was reported *in vitro* in isolated phycobiliproteins ^37^. It was related to photoisomerization of the billins into a form more prone to quenching, due to interactions of the solvent with the chromophores that are influenced by temperature. Maksimov and co-workers reasoned that the interaction between monomeric subunits of phycobiliprotein trimers prevents photoisomerization of the chromophores. Once this interaction is weakened due to the decrease in temperature, photoisomerization takes place. In our case the measurement was done *in vivo,* where linker proteins occupy the center of the trimer. Nevertheless, we were able to detect this anomaly in intact cell samples. The *in vivo* temperature anomaly had a different trend in medium and low light cells. While in medium light cells the rise began immediately upon cooling, low light grown cells maintained a constant lifetime down to lower temperatures (to about 280K), and their rise in lifetime is more gradual. According to the proposed explanation for the rise in lifetime, this difference indicates stronger interaction between the chromophores in the low light grown PBS, supporting our hypothesis.

Fig. 6B and fig. 6C show the emission spectra at a few chosen temperatures as a function of temperature. In both medium and low light cells, emission intensity increases dramatically with the decrease in temperature, as expected when decreasing the heat dissipation component. The main difference was the change in the ratio between the PUB emission (560 nm) to PEB emission (574 nm). At room temperature, the emission of both pigments comprised one visible peak due to thermal broadening. At lower temperatures, the two spectral lines narrowed and the emissions of the two pigments were resolved. While under low light the PUB:PEB emission peaks ratio decreases significantly with temperature (down to 0.45 at 5K), under medium light the ratio is closer to 1 (reaching 0.8 at 5K). Since the PUB and PEB content remained constant while cooling down, it implies that under low light a larger fraction of the energy absorbed by PUB was transferred to PEB and emitted from PEB, even at 5K, proving stronger coupling between the PUB-PEB pigments. It is interesting to note that the PBS under low light has a higher PUB:PEB ratio (as inferred from fig. 3 and illustrated in fig. 4). In other words, even though the PUB content under low light is higher, we observe less PUB emission than in medium light. These results further support the hypothesis that in cells acclimated to low light, the coupling between the chromophores in the PBS is stronger. The stronger coupling leads to the higher energy transfer rate.

## Conclusions

Taking advantage of the extensive photo-acclimation capacity of *Synechococcus* WH8102, we were able to study physical mechanisms governing the acclimation of the photosynthetic apparatus. Cells displayed extreme morphological flexibility, acclimating to the irradiance regime within days. When experiencing low blue light, growth rates were slower, cells were bigger and contained more thylakoid membranes. The number of photosynthetic units increased, but what we find most fascinating is that each of these units was more efficient under a low light regime.

The PBS was bigger under low light, with longer rods containing additional PUB pigments which increase the absorption cross-section at 495 nm. Surprisingly, instead of decreasing the rate of energy transfer, as was reported in plant systems ^35^, the longer rods exhibit faster energy transfer, as seen by the shorter lifetime measured as a function of temperature. Several lines of evidence support the hypothesis that improved energy transfer is achieved by stronger coupling between the pigments. Coupling strength according to the semi-classical FRET theory is a function of dipole strength, distances, orientation and energetic overlap between the pigments in the exciton transfer network. Increased efficiency over a larger pigment network suggests the involvement of coupling that is stronger than predicated by FRET calculations. While it is too early to pinpoint the exact energy transfer mechanism, the photo-acclimation response studied here is a step towards a better understanding of these processes. Unveiling the mechanism by which cyanobacteria tune energy transfer will help understand photo-acclimation, and may pave the way to bio-inspired nano-scale energy-transducing devices with a broad dynamic range.

## Materials and Methods

### Growth conditions and cellular morphology measurements

#### Synechoccocus

WH8102 cultures were grown in ASW medium under continuous blue light (spectrum in Supplementary fig.1), in 10 and 150 *µmol photons m*^−2^*s*^−1^, referred to as “low light” and “medium light” respectively. For FACS analysis, one ml samples were collected every second day, preserved in 0.2% glutaraldehyde grade II (Sigma), frozen in liquid nitrogen and stored at –80°C. Cells were counted using FACS Aria III flow cytometer-sorter (BD Biosciences, San Jose, CA) using 488nm laser. The flow rate was 14 μl min^−1^, data was collected for 1 min. Confocal microscopy was preformed using an Olympus FV-1200 confocal microscope Olympus, Japan. For transmission electron microscopy (TEM), cells were harvested by centrifugation at 8000g for 5 min, washed twice in ASW and chemically fixed in 2% paraformaldehyde and 2.5% Glutaraldeyde in 0.1M Cacodylate buffer (pH 7.4). Sample preparation followed standard procedures. Sections were sequentially stained with Uranyl acetate and Lead citrate for 3 minutes each and viewed with Tecnai 12 TEM 100kV (Phillips, Eindhoven, the Netherlands) equipped with MegaView II CCD camera.

### Physiological parameters, pigment and protein composition

Oxygen Evolution was measured using Clark-type electrode (Hansatech Instruments Ltd, King’s Lynn, Norfold, UK). Measurements were performed using the same blue LED illumination under which the sample grew. Light intensities were altered by changing the distance of the light source from the culture and by using ND filters for attenuation. Chlorophyll fluorescence kinetic parameters were measured using a PAM 2500 (Walz, Effeltrich, Germany). Fm was measured following the addition of DCMU under actinic light.

Hydrophobic pigments were extracted in 100% methanol and the pellet containing hydrophilic phycobilins was suspended in TMN.

### Spectroscopic measurements

Steady state Absorption and Fluorescence were performed within an integrated sphere to reduce scattering, using a Horiba PTI Quantamaster (Kyoto, Japan).

Temperature dependent time-resolved fluorescence measurements were carried out using a time-correlated single-photon counting (TCSPC) setup built on attoDRY800 cryo-optical table. Excitation was done with a Finaium WhiteLase SC-400 supercontinuum laser monochromatized at 485 nm at a repetition rate of 40MHz. Detected with MPD PD-100-CTE-FC photon counter and PicoHarp300. The samples were trapped between two sapphire windows with a Teflon O-ring in between, in their original medium with no additives. This sample holder was put inside a vacuum chamber, thermally coupled to a cold plate that is gradually cooled down to 4 k. Sample temperature was measured along the process with a Cernox DT-670-SD attached to the sapphire window. There may be an offset of up to a few degrees between the temperature recorded by the sensor and the actual temperature of the sample. Lifetime data was fitted using DecayFit 1.4 software and a three exponential decay model.

Temperature dependent emission spectra were recorded at the same cryogenic setup: excited with a CPS450nm laser diode and emission detected with Flame spectrometer by Oceanview.

## Supplementary Information

**Fig. S1.**
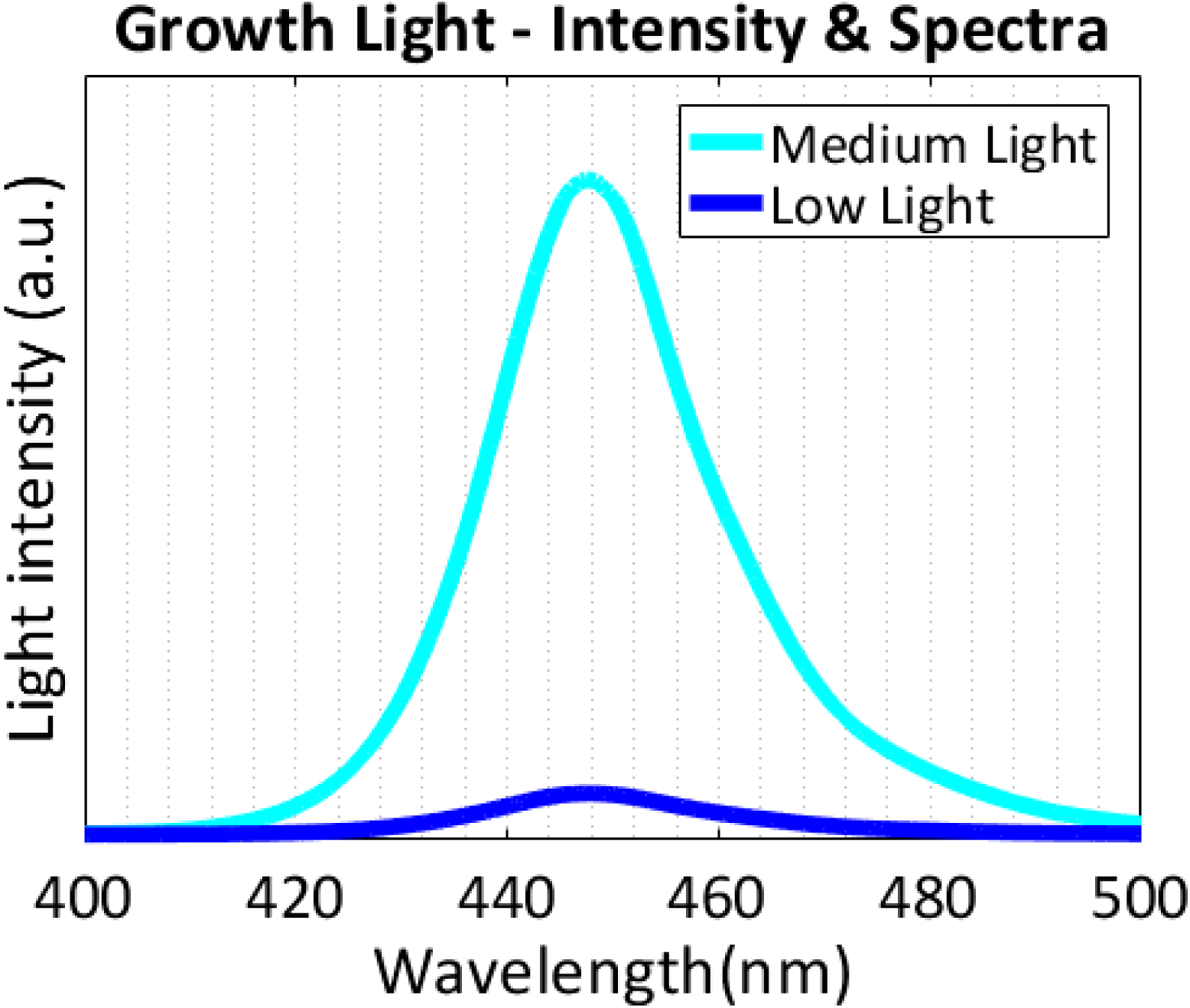
Spectra of LED illumination under which the bacteria cultures were grown. Medium light is 150 and low light is 10 ***µmol photons m*^−2^*s*^−1^**. The spectra resembles the spectra found within the water column.

**Fig. S2.**
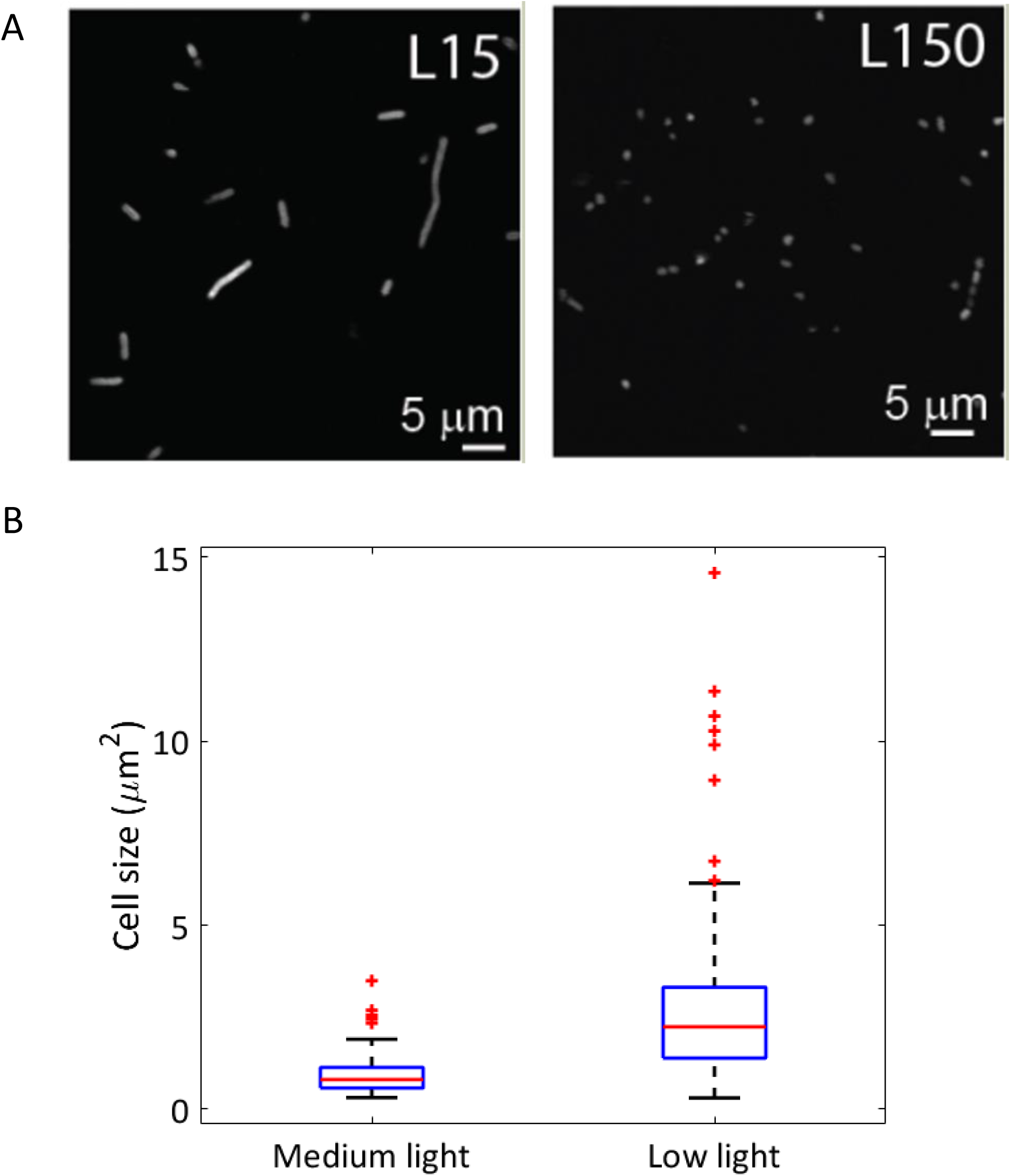
(A) Fluorescence confocal imaging of low light (L15) and medium light (L150) grown cells. (B) Quantification of cell sizes from confocal microscopy images. The box plot central mark indicates the median, the bottom and top edges of the box indicate the 25th and 75th percentiles. The whiskers extend to the most extreme data points not considered outliers, shown as red pluses.

**Fig. S3.**
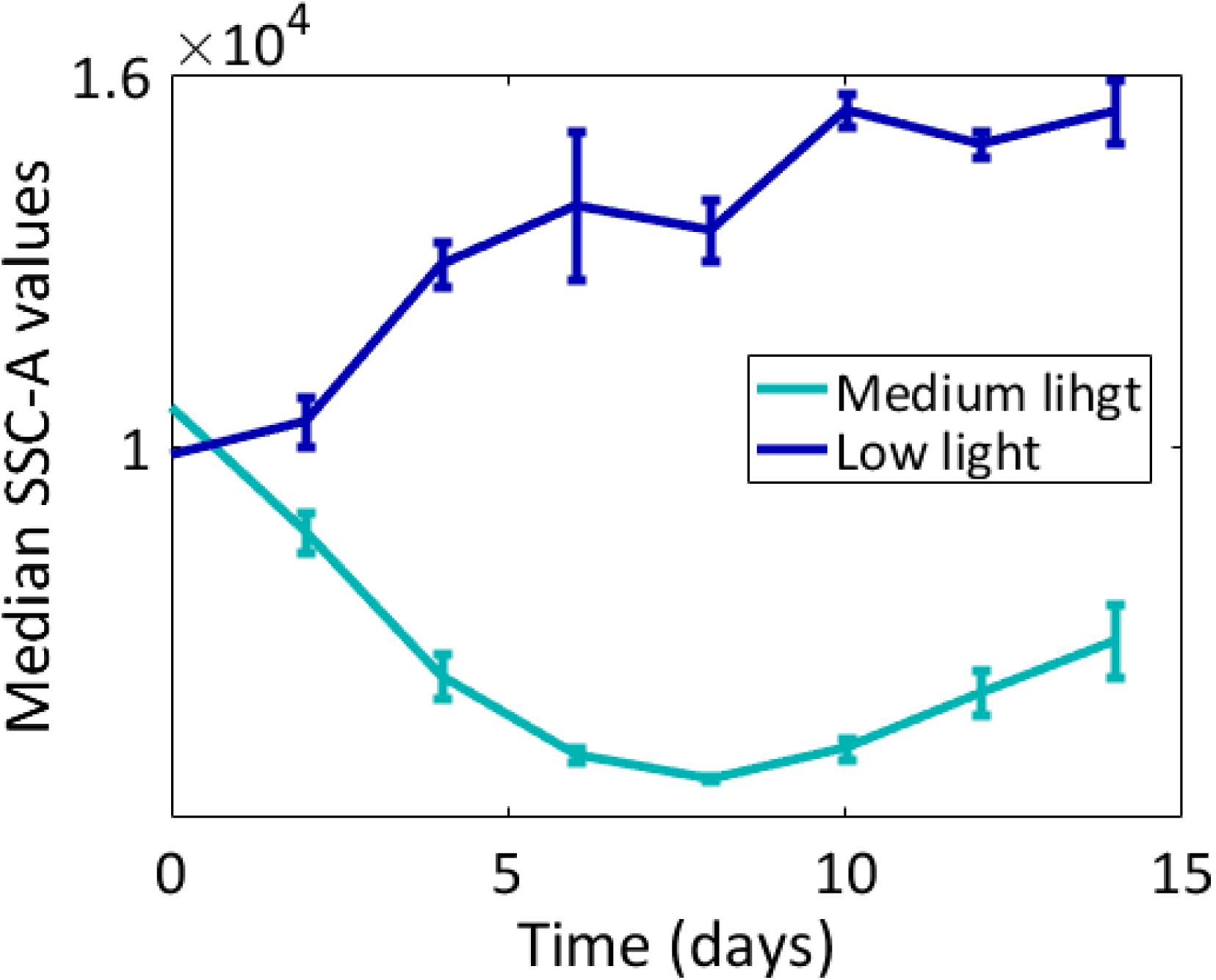
Progression of changes in SSC-A Median values. Averages and SD were derived from three independent cultures.

**Fig S4.**
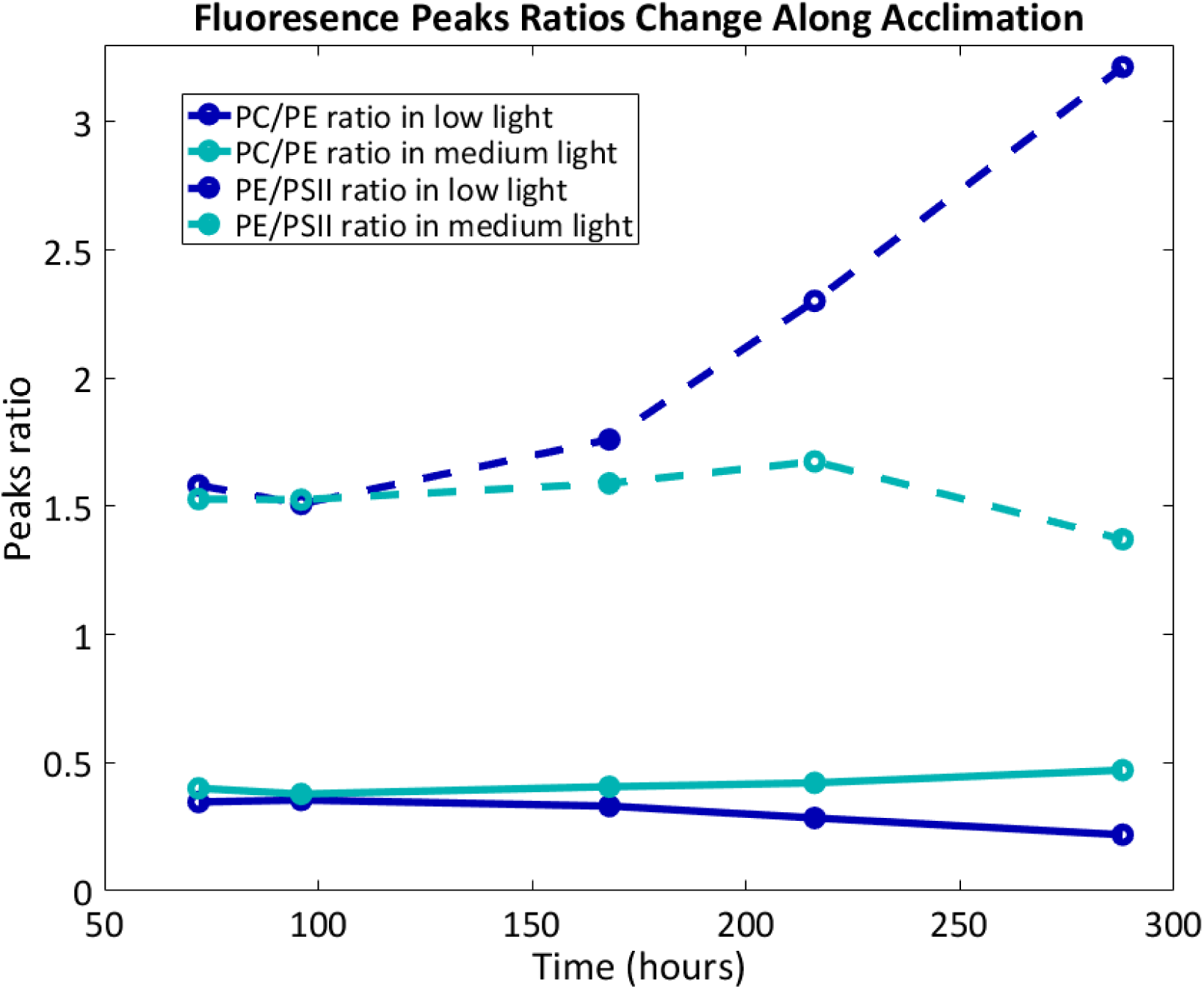
Fluorescence spectra following excitation at 497 nm were recorded along the acclimation process. The ratios between the different pigments’ peaks change along the process. Cultures were treated with 5 mM DCMU prior to measurement to allow better comparison of PSII emission.

**Fig S5.**
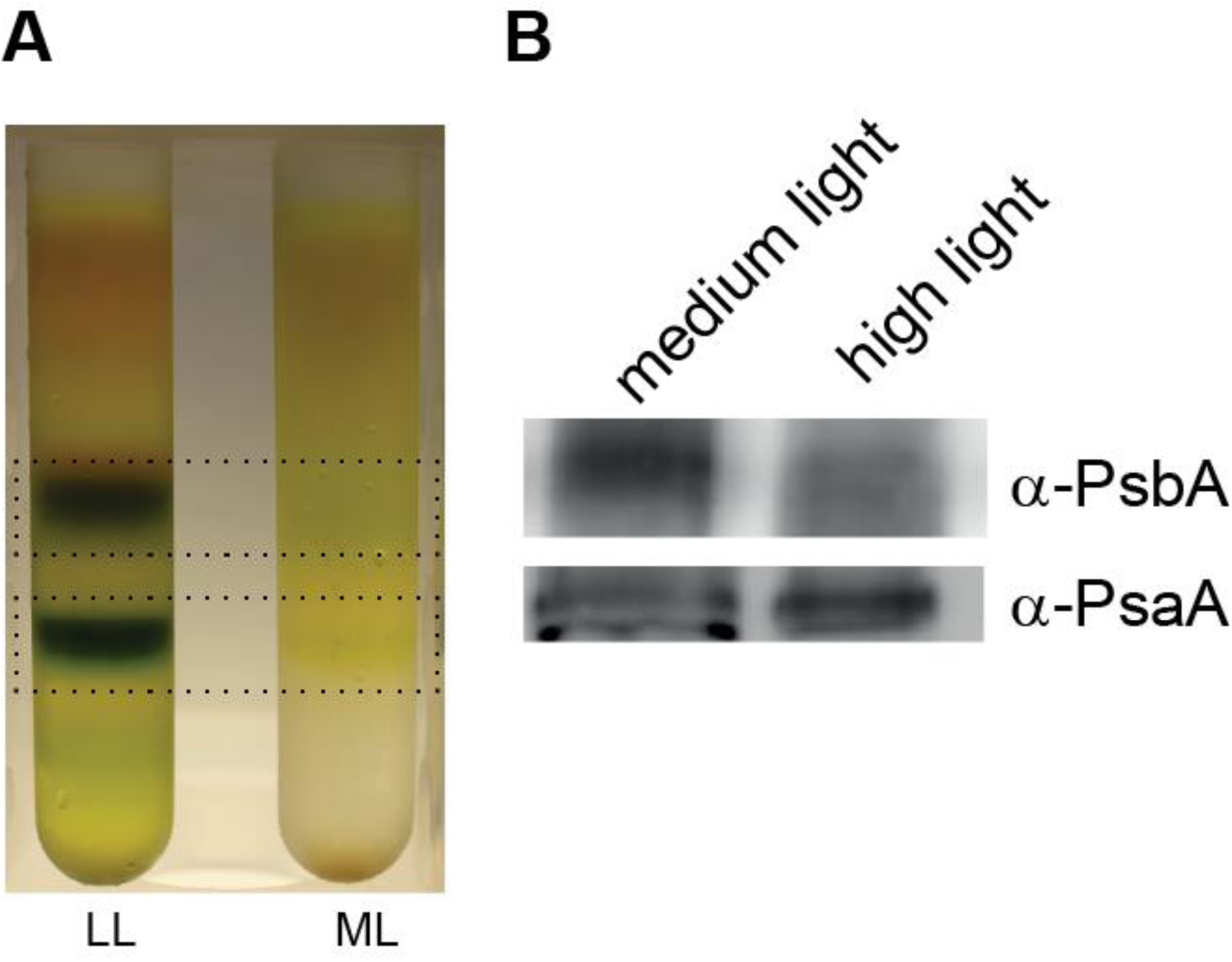
(A) Thylakoids preparation and photosystem complexes separation. Following 7 days incubation under medium or low blue light, cells were centrifuged (12,800g for 7min) and washed twice with buffer A (50mM Tris HCl pH8; 10mM NaCl; 1mM EDTA; 200mM Sorbitol). Finally, cells were resuspended in 5 ml buffer A with the addition of anti-protease cocktail (Roche). Cells were broken using French Press, centrifuge for 3 min using 1,160g in 4C. The supernatant was collected to a new test tube and was centrifuge for 1hr using 95,000 g in 4 oC. The pellet containing thylakoids, was resuspended in small volume of buffer A. Proteins concentration was determined using Bradford reagents. Solubilization of membrane pigment-protein complexes was done using dodecyl-maltoside (DDM 1.5 % per 1 mg protein). Thylakoids were incubated than for 30 min in ice. Following incubation, thylakoids were centrifuged for 10 min using 17,000g in 4C. The supernatant was loaded on sucrose gradient (0-40% in buffer A with the addition of 0.05% DDM). Gradients were centrifuged 16 h at 36,000 rpm (SW41) in 4C. (B) Quantification of PsaA and PsbA in the total membrane fraction by SDS-PAGE and immunoblotting. The data indicates a slightly higher PSI/PSII protein content under medium light.

**Figure S6.**
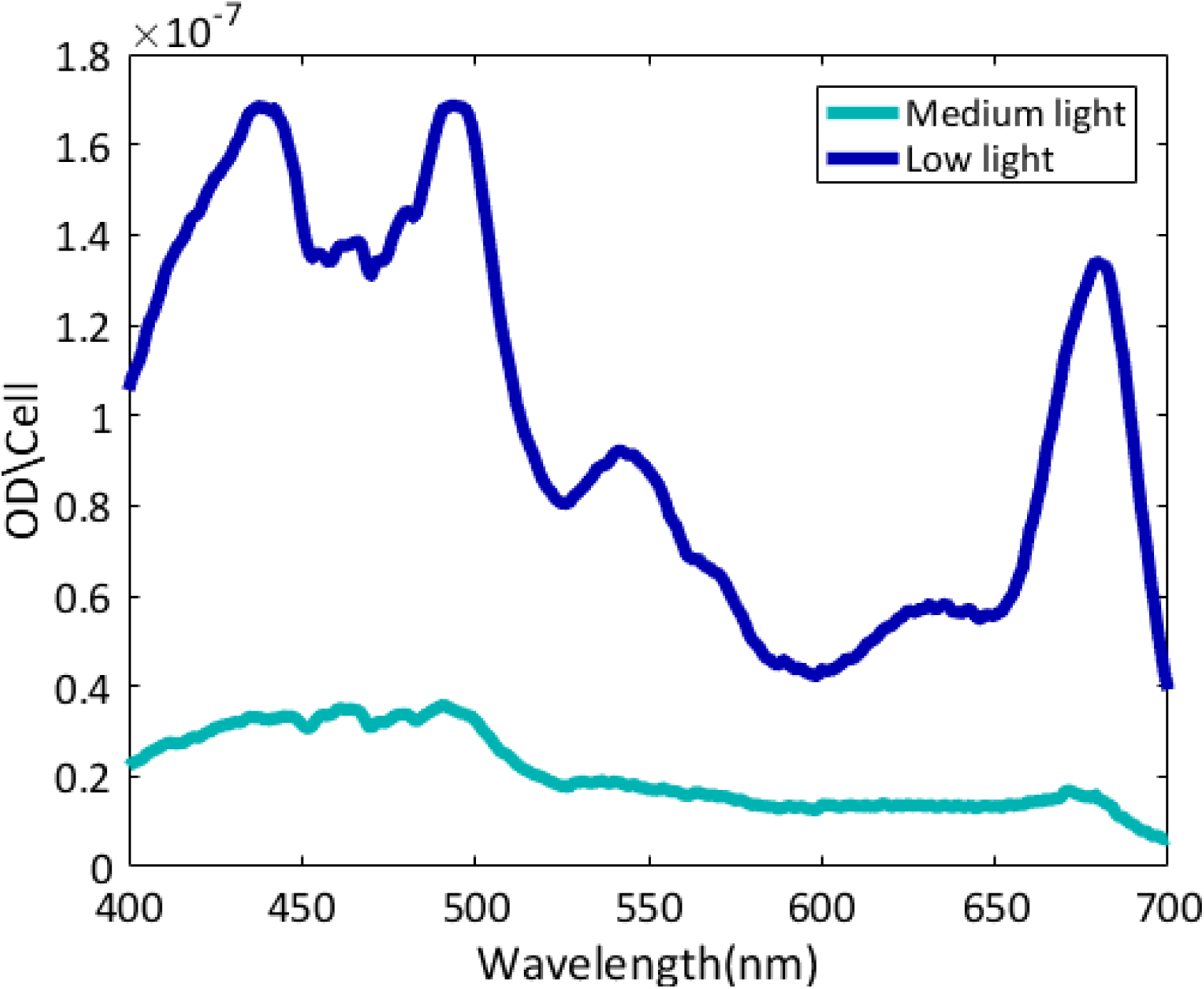
Absorption spectra of medium light and low light grown cells *in vivo,* taken prior to fractionation of the cells and extraction of the pigments.

## Footnotes

1 Aristotle, Metaphysics, book VIII part 6 (350 B.C.E).

